# Tractable and Thermodynamically Consistent p*K*_a_ Prediction via First-Dissociation Aggregation

**DOI:** 10.64898/2026.09.30.755586

**Authors:** Zhuoyan Liu, Qiuyin Zhu, Qingkun Li, YuanXi Zhu, You Xu, Bin Dai, Yongsheng Li, Zhenming Liu, Shengwen Yang

**Affiliations:** Key Laboratory of Xinjiang Endemic Phytomedicine Resources, Ministry of Education; School of Pharmacy, Shihezi University, Shihezi 832002, China; School of Chemistry and Chemical Engineering/State Key Laboratory Incubation Base for Green Processing of Chemical Engineering, Shihezi University, Shihezi, 832003, China; Lab of Low-Dimensional Materials Chemistry, Key Laboratory for Ultrafine Materials of Ministry of Education, Frontier Science Center of the Materials Biology and Dynamic Chemistry, Shanghai Engineering Research Center of Hierarchical Nanomaterials, School of Materials Science and Engineering, East China University of Science and Technology, Shanghai 200237, China; State Key Laboratory of Natural and Biomimetic Drugs, School of Pharmaceutical Sciences, Peking University, Beijing, 100191, China; Beijing Advanced Center of Cellular Homeostasis and Aging-Related Diseases, Institute of Advanced Clinical Medicine, Peking University, Beijing, 100191, China

**Keywords:** p*K*_a_ prediction, protonation equilibria, thermodynamics,graph neural networks, machine learning

## Abstract

Ionization governs molecular behavior, yet predicting aqueous pKa accurately and tractably remains a fundamental challenge. Rigorous ensemble methods require enumerating a protonation-state space that grows exponentially with the number of ionizable sites, while fast graph predictors return numbers without thermodynamic consistency. Here we derive an exact thermodynamic identity showing that the macroscopic first-dissociation constant is fully determined by protonated-microstate populations and unique first-deprotonation events, eliminating the deprotonated ensemble algebraically and reducing aggregation to O(n+E) operations. We implement this identity in DTi-pKa, a dual-head graph neural network whose free-energy and dissociation heads are coupled by thermodynamic constraints. On four external benchmarks (355 molecules), DTi-pKa achieves a pooled MAE of 0.5665 pKa units, with 0.6285 on the 169 records lacking training-cache matches, while providing site-resolved micro-pKa values and protonation-state populations. Controlled ablations show that constraint placement at the level of the reported macroscopic equilibrium matters more than microscopic label precision—a design principle transferable to physics-informed machine learning beyond chemistry. Closure diagnostics expose numerical self-consistency as an open frontier, which we report transparently. By replacing an intractable computation with a physical identity, DTi-pKa unifies macroscopic accuracy and microscopic interpretability in a single framework.

## Introduction

Embedding physical laws into machine learning systems is a central challenge in AI for Science^1,2^. Physics-informed neural networks have improved prediction wherever first principles are known^2,3^, yet most molecular graph models remain purely data-driven: they interpolate within training distributions and offer no guarantee of thermodynamic consistency^4,5^. A particularly severe case is solution-phase ionization equilibria, where rigorous ensemble methods must enumerate a protonation-state space that grows as 2^N^ with the number of ionizable sites. Fast graph predictors avoid this cost but return numbers without an equilibrium, and their macroscopic and site-level outputs are not guaranteed to agree for multi-site molecules. This trade-off—thermodynamic rigor versus computational tractability—has confined physics-grounded methods to candidate-level studies and kept screening-scale predictors physically unconstrained.

Charge is the most consequential single attribute of a drug molecule in water. Approximately 70% of FDA-approved oral drugs carry at least one ionizable group^6,7^. The ionization state at physiological pH, encoded by pKa, governs aqueous solubility, permeability, protein binding, metabolic stability and off-target toxicity, the properties that decide ADMET^8,9,10^. A pKa error of one unit shifts the ionized-to-neutral ratio tenfold, enough to move a lead compound across the accept-or-reject boundary in early discovery^10^. Yet pKa is not a local property of one functional group. In polyfunctional molecules, proton transfer, tautomerism and charge redistribution couple the ionizable sites, so every measured macroscopic pKa is an equilibrium over an ensemble of protonation microstates^11,12^. The experiments that resolve such equilibria (potentiometry, spectrophotometry, capillary electrophoresis) remain slow, costly and out of reach at screening-library scale^13^. Ionizable molecular properties now drive machine-learning-guided delivery design at library scale^14^

Computational prediction is therefore the practical route to ionization profiling, and two families dominate. Fragment-based and graph-neural predictors^15-18^ are fast enough for screening and accurate on familiar chemotypes. They regress pKa as a standalone endpoint, however: they return a number, not an equilibrium, and their macroscopic and site-level outputs are not guaranteed to agree for multi-site molecules. Recent refinements push accuracy with quantum-derived features^19,20^, sequence-based unification^21^ and thermodynamics-informed networks that report strong blind-prediction results^4,22^. Physics-grounded methods construct the equilibrium explicitly^22^. Constant-pH molecular dynamics^23^ and free-energy methods such as Epik^24,25^ and MCCE2^26^ integrate microscopic-state thermodynamics and are regarded as reference-grade, but cost hours per molecule. Machine-learning methods that borrow this rigor inherit its price. Uni-pKa builds free-energy frameworks whose partition functions still enumerate large protonation ensembles for each molecule^22^. QupKake attains accuracy through costly quantum-chemical descriptors, which limits its scalability^19^. TripKa trains against quantum-level targets^20^. In every thermodynamics-aware method the number of conceivable protonation states grows as 2^N^ with the number of ionizable sites N. This protonation-state enumeration explosion, not accuracy, is what keeps rigorous prediction out of screening pipelines.

Two challenges follow. First, no single framework delivers macroscopic accuracy and microscopic interpretability together; models optimize one at the expense of the other. Second, the methods that respect thermodynamics remain trapped behind the 2^N^ enumeration wall. By thermodynamically consistent we mean that predicted microscopic dissociation constants and molecular free energies jointly satisfy the identity governing the macroscopic equilibrium. Most black-box regression models do not enforce this property. A broader development suggests a way forward: physics-informed neural networks, which embed governing identities as constraints on learning, have improved prediction wherever first principles are known^2,3,4^. The present work asks whether such an identity can do something stronger than regularize, whether it can replace the intractable computation itself.

The key observation is that it can. The macroscopic first-dissociation constant is the ratio of total product activity to total reactant activity. The products of first dissociation are nothing other than the protonated microstates with one proton removed, and the microscopic dissociation constants relate each product’s activity to its parent’s. Substituting these relations into the macroscopic definition, the deprotonated-state activities cancel algebraically. The macroscopic pKa is exactly a population-weighted sum over the protonated microstates and their unique first-deprotonation events, a linkage relation in the spirit of classical binding-polynomial analysis^27^. Aggregation then requires O(n+E) operations for n retained protonated microstates and E unique first-dissociation paths. The deprotonated ensemble, the exponential part, need never be enumerated.

We implement this identity in DTi-pKa, a dual-head graph neural network whose free-energy and dissociation heads are coupled by thermodynamic training constraints. The design principle is transferable: embed an identity that eliminates an intractable computation, enforce it as a soft constraint, and measure with ablations and closure diagnostics whether the constraint is causal rather than decorative. We test the framework as a claim about mechanism rather than correlation. Controlled ablations isolate each constraint, and closure diagnostics measure how far the trained heads satisfy the identity. A generalization audit separates memorization from extrapolation, and failure cases localize where the method’s assumptions break. All evaluations run on four external benchmarks (SAMPL6, SAMPL7, SAMPL8 and Novartis) that were never touched during model selection^28–32^. By replacing an intractable computation with a physical identity, DTi-pKa unifies macroscopic accuracy and microscopic interpretability in a single framework, and establishes a design principle with reach beyond pKa prediction.

## Results and Discussion

### An exact thermodynamic identity eliminates the protonation-state enumeration barrier

DTi-pKa begins not from a regression target but from the equilibrium itself. Consider n protonated microstates HA_i_ and the distinct products A_alpha_ ^-^ reachable by removing one proton from them. Under an ideal-solution standard-state convention with dimensionless activities, the macroscopic first-dissociation constant is

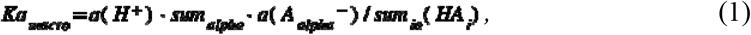

where a(X) denotes the activity of species X. The standard state fixes a(H^+^) = 1; it does not prescribe a physical proton concentration of 1 mol/L. We write the dimensionless protonated-state free energy *g*_*i*_ and the microscopic log-equilibrium constant *ell*_*i,alpha*_ as

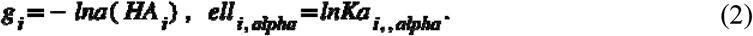

With a(H^+^) = 1, mass action gives *Ka*_*i,alpha*_ = exp(*ell*_*i,alpha*_) and *pK*_*i,alpha*_ = -*ell*_*i,alpha*_/ln 10. This convention fixes the sign explicitly and prevents a conventional reaction free energy from being inserted with the wrong exponential sign.

Interconversion among protonated microstates obeys *Ka*_*ik*_ *= a(HA*_*k*_*)/a(HA*_*i*_*) = exp(g*_*i*_ *-g*_*k*_). The equilibrium population of HA_i_ within the protonated ensemble is

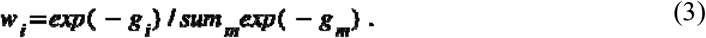

For each distinct product *A*_*alpha*_^*-*^, one thermodynamically equivalent generating path is retained; let *U*_*i*_ denote the unique products assigned to *HA*_*i*_. This partition prevents double counting when different sites or protonated microstates converge on the same product. Substituting the microscopic mass-action relations into equation (1) eliminates every deprotonated-state activity and gives the central identity

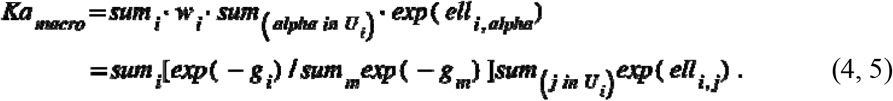

Equation (4) is an algebraic identity for the specified state set, not an approximation: its exactness requires dimensionless activities, inclusion of the relevant protonated microstates and distinct first-dissociation products, and consistency among convergent paths. Product activities are eliminated algebraically; they are not assumed to be absent. Because the model predicts *ell*_*i*_,_*alpha*_ directly, independent product free energies do not enter the macroscopic sum. Canonical first-dissociation products are nevertheless generated for unique-path bookkeeping and for the teacher and diagnostic relations.

Substituting the Boltzmann-weighted population *w*_*i*_ from equation (3) and taking the negative logarithm gives a numerically stable form. Each parent contribution is evaluated by log-sum-exp with maximum subtraction:

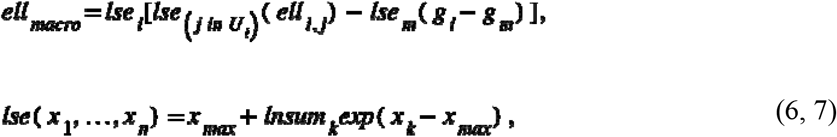

with the terms sorted in descending order and *pKa*_*macro*_ *= -ell*_*macro*_*/ln 10*. The aggregation requires O(n+E) operations for n retained protonated microstates and E unique first-dissociation paths. This statement concerns aggregation, not the separate generation of protomers and tautomers.

The identity removes the combinatorial barrier at the level of the physics, not by truncation or approximation. Figure 2b shows that the path count grows polynomially where the full binary state space grows exponentially. The barrier that remains is the enumeration of protonated microstates, which is bounded by the number of ionizable sites times the chemically accessible tautomers of each. A 110-molecule audit on a consumer GPU gave median total inference times of 187, 575 and 2,554 ms for one, two and three-or-more detected sites (Fig. 2c, Supplementary Fig. S4 and Table S7). These measurements establish practical tractability for the tested implementation, not an empirical asymptotic law. Figure 2a makes the population-weighting principle concrete on a molecule with three titratable sites, whose two dissociation paths converge on the same product and are counted once.

**Fig. 1.**
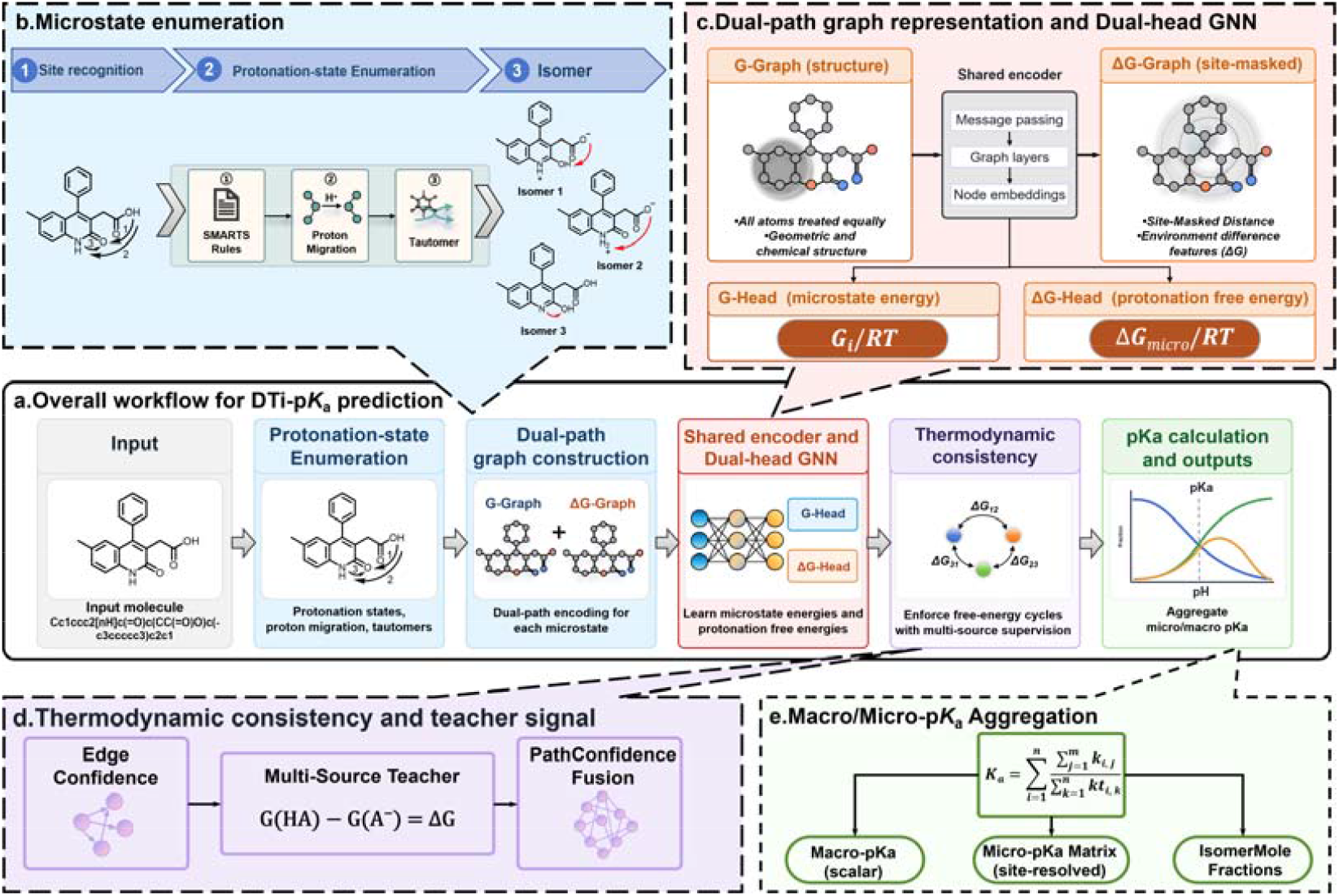
Overview of DTi-pKa. **a**, Overall workflow for DTi-pKa prediction: input molecules pass through site recognition and microstate enumeration. **b**, Microstate enumeration and Dual-path graph featurization. The G-graph treats all atoms equally; the ΔG-graph uses site-masked distances to encode the dissociation site. **c**, Dual-head graph neural network (GNN). A shared graph encoder feeds two heads: a molecular free-energy (G) head predicting a dimensionless scalar G/RT, and a site- and path-resolved dissociation head (ΔG-head) predicting ell = ln Ka, stored as ΔG_micro/RT. **d**, Thermodynamic closure. Confidence-weighted multi-source teachers connect the two heads during training via same-product and fanout relations, G(HA) = G(A□) + ΔG and G(A □) = G(HA) − ΔG, with path confidence from a PathConfidence network. **e**, Macro/micro-pKa aggregation. Unique first-dissociation paths are aggregated using the exact identity 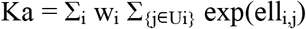, where w_i_ are Boltzmann-weighted protonated-microstate populations and U_i_ are unique products assigned to parent state i. The deprotonated ensemble cancels algebraically, reducing aggregation to O(n+E) operations for n retained protonated microstates and E unique first-dissociation paths. Outputs are macroscopic pKa (scalar), site-resolved micro-pKa matrix, and isomer mole fractions. Color legend: blue, G-head; orange, ΔG-head; purple, shared encoder; green, aggregation. SMARTS, Smiles ARbitrary Target Specification; RT, thermal energy.

**Fig. 2.**
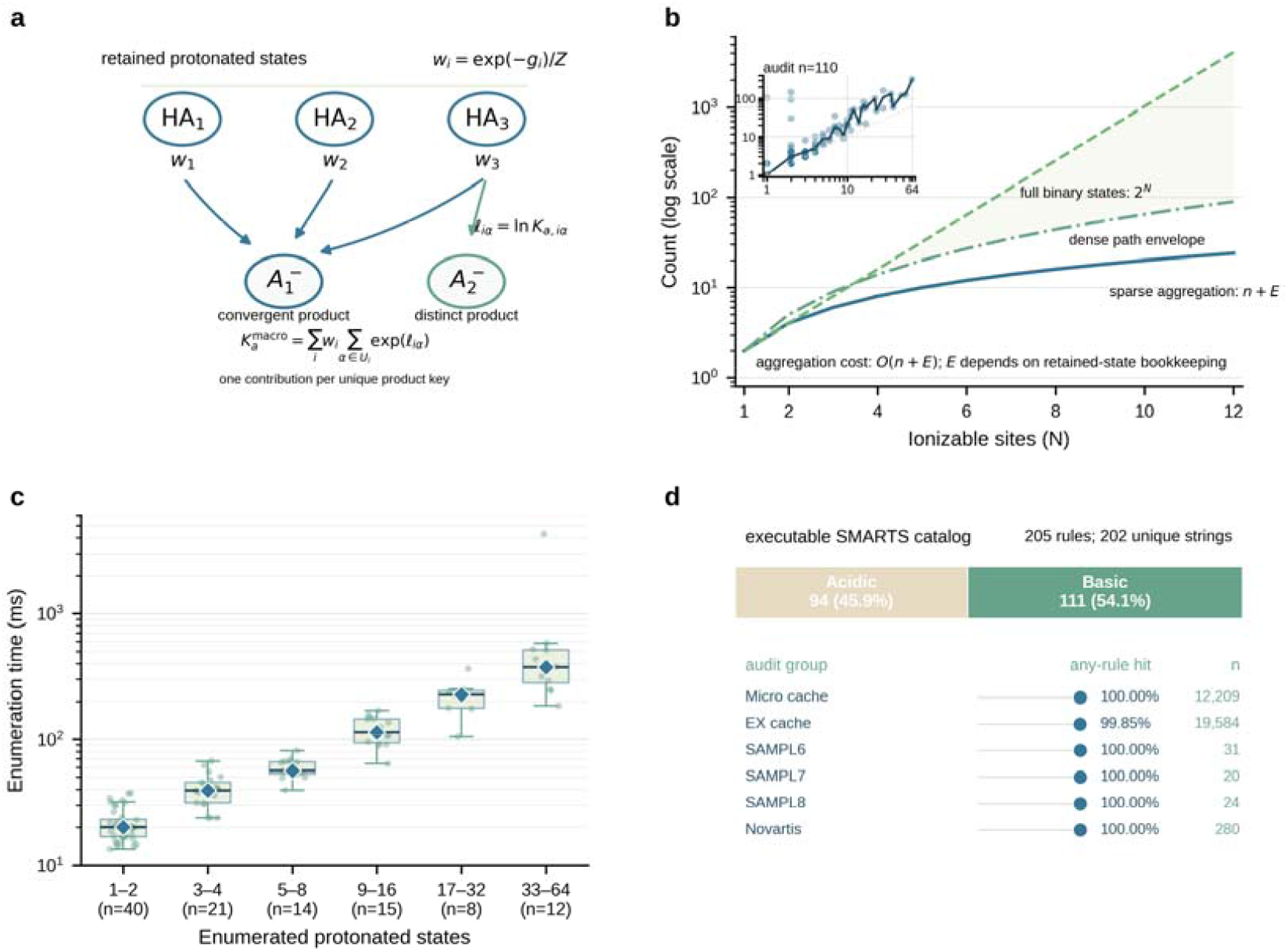
Thermodynamic basis and computational scaling of first-dissociation aggregation. **a**, Population-weighted first dissociation for alternative protonated microstates. The macroscopic constant is 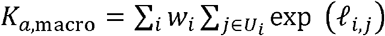 where *w_i_* is the Boltzmann-weighted population of protonated microstate HA_*i*_ and *U_i_* contains the unique first-dissociation products assigned to HA_*i*_ . Convergent paths to the same product are counted once. **b**, Theoretical number of unique first-dissociation paths (sparse aggregation, *O*(*n*+*E*)) compared with the full binary protonation-state space (2^*N*^) as a function of the number of ionizable sites N. Note the logarithmic scale. **c**, CPU enumeration latency for representative molecules. Box plots show the distribution of enumeration times (ms) grouped by the number of enumerated protonated states; diamonds indicate medians; the audit included 110 molecules. **d**, Audit of the executable SMARTS catalogue (205 rules, 202 unique strings; 94 acidic labels, 111 basic labels). Bars show the fraction of records with any-rule hits for each dataset; numbers above bars indicate the number of records (n). Agreement with supplied acidic-site labels is reported only where such labels exist and is not a formal precision/recall metric.

### External benchmark performance of DTi-pKa

The publication checkpoint was selected by validation macro-pKa mean squared error (MSE 0.4563) before any external benchmark analysis, and was then evaluated once on SAMPL6^28,29^, SAMPL7^30^, SAMPL8^31^ and Novartis^32^. All 355 external records produced valid predictions (Fig. 3). DTi-pKa attained a pooled mean absolute error (MAE) of 0.5665 pKa units, a root-mean-square error (RMSE) of 0.7227 and a coefficient of determination (R2) of 0.9101 (Supplementary Table S4). Two qualifications materially affect this estimate. First, canonical-SMILES auditing shows that 186 of the 355 records (52.4%) exactly match the retained training cache. On the 169 records without such matches the metrics are MAE 0.6285, RMSE 0.8027 and R2 0.8674 (Supplementary Table S4). Second, the pooled value averages two record types: site-resolved (Micro) records (n = 150, MAE 0.5027) and macroscopic (EX) records (n = 205, MAE 0.6132). These are different estimands and are reported separately throughout.

**Fig. 3.**
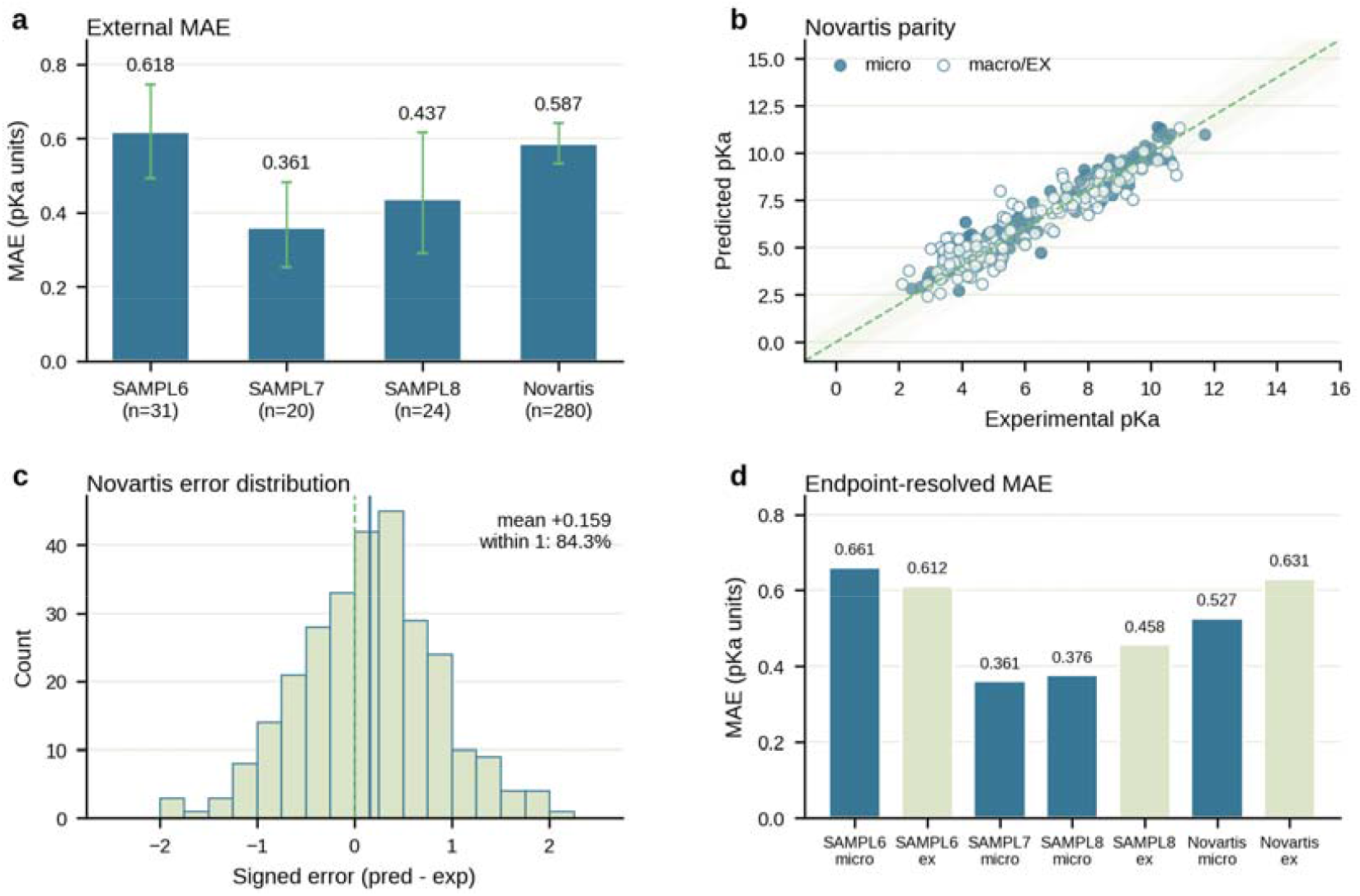
External benchmark performance of DTi-pKa. **a**, Mean absolute error (MAE) on four external benchmarks (SAMPL6, SAMPL7, SAMPL8, Novartis; n = 355 total). Error bars represent 95% nonparametric bootstrap confidence intervals computed from matched per-record prediction errors (10,000 resamples). **b**, Parity plot for Novartis (n = 280). Filled symbols denote site-resolved (Micro) records; open symbols denote macroscopic (EX) records. The diagonal line indicates perfect agreement; shaded bands indicate ±1 and ±2 pKa units from the line of identity. **c**, Distribution of signed errors (predicted − experimental) for Novartis; the mean signed error is +0.159 pKa units. **d**, Endpoint-resolved MAE by benchmark, shown separately for Micro and EX records. Error bars, where present, represent standard errors.

Performance varied by benchmark. SAMPL7 was the strongest set (MAE 0.3607, RMSE 0.4473, R2 0.9637). All 20 of its records are exact cache matches, so it is reported as a cache-matched set rather than as evidence of generalization. SAMPL6 carried the largest error tail (MAE 0.6181, RMSE 0.7168, R2 0.9291), and SAMPL8 and Novartis were intermediate (MAE 0.4372 and 0.5866). On the macroscopic SAMPL8 stratum, DTi-pKa’s MAE (0.4575) is similar to the reported H-SPOC value (0.449)^33^. On SAMPL6, DTi-pKa trails H-SPOC and QupKake^19^ on the reported RMSE. On Novartis, DTi-pKa has RMSE 0.7487 overall (0.8091 on the macroscopic stratum), against the locally recorded Uni-pKa^22^, TripKa^20^ and QupKake^19^ values of 0.840, 0.802 and 0.790. These are literature-context comparisons rather than controlled head-to-head experiments, because training data, endpoint definitions, state generation and evaluation protocols differ, and the published values carry no uncertainty (Supplementary Table S5). The standard error of our per-benchmark MAE is 0.0663 (SAMPL6), 0.0607 (SAMPL7), 0.0845 (SAMPL8) and 0.0279 (Novartis), so differences below roughly one standard error are not interpretable. Rather than treating the SAMPL6 deficit as a scoreboard entry, we took it as the decisive question for the failure analysis below. Which feature of that chemical space breaks the assumptions of the aggregation, and which does not?

### Generalization audit: memorization versus extrapolation

Canonical-SMILES auditing identified 186 exact overlaps between the 355 external records and the retained training cache (Supplementary Information S10). They fall as 12 in SAMPL6, 20 in SAMPL7, 7 in SAMPL8 and 147 in Novartis. The overlap is concentrated in the site-resolved records: all 150 Micro records are exact matches, against 36 of 205 EX records. One record (Novartis_135_) is flagged as non-overlapping although its nearest cache neighbour has Tanimoto similarity 1.0; the stored flag is retained as approximate. The audit covers the fine-tuning cache only, and overlap with the roughly 1.99-million-row ChEMBL pretraining corpus was not audited.

Similarity-stratified results sharpen the picture. The combined MAE was 0.5934 for 44 records with nearest-neighbour Tanimoto similarity below 0.40 and 0.6636 for 86 records in the 0.40-0.59 interval. It was 0.5971 for 37 records in the 0.60-0.79 interval and 0.5098 for 188 records at or above 0.80 (Supplementary Information S10). The high-similarity group contains 186 exact overlaps and should not be interpreted as a clean generalization stratum. The low-similarity subset remains a useful audit of performance away from exact cache matches, but its size and benchmark composition do not establish broad chemical-space generalization. The benchmark sets were not used for checkpoint or hyperparameter selection. The pooled results are nevertheless reported alongside the non-overlap metrics (Supplementary Table S4), so that memorization and extrapolation remain visible side by side.

### Constraint placement at the macroscopic level improves generalization more than microscopic label precision

To attribute external performance to individual mechanisms, we trained six controlled switch-off variants (A1-A6) in which only the target component was removed (Fig. 4d, Supplementary Table S6 and Information S9). Removing any single component consistently increased the pooled external MAE. The paired delta-MAE values range from +0.0449 to +0.0760 pKa units, with 95% bootstrap intervals that exclude zero at the pooled level. These variants remain single-seed runs whose logged durations were not recorded, and the intervals are unadjusted for multiple comparisons. The per-benchmark A6 intervals include zero on SAMPL6 and SAMPL8. The results therefore support component-associated differences under the recorded protocols rather than definitive causal attribution. With that discipline stated, the ranking of the degradations traces the flow of thermodynamic information through the model.

**Fig. 4.**
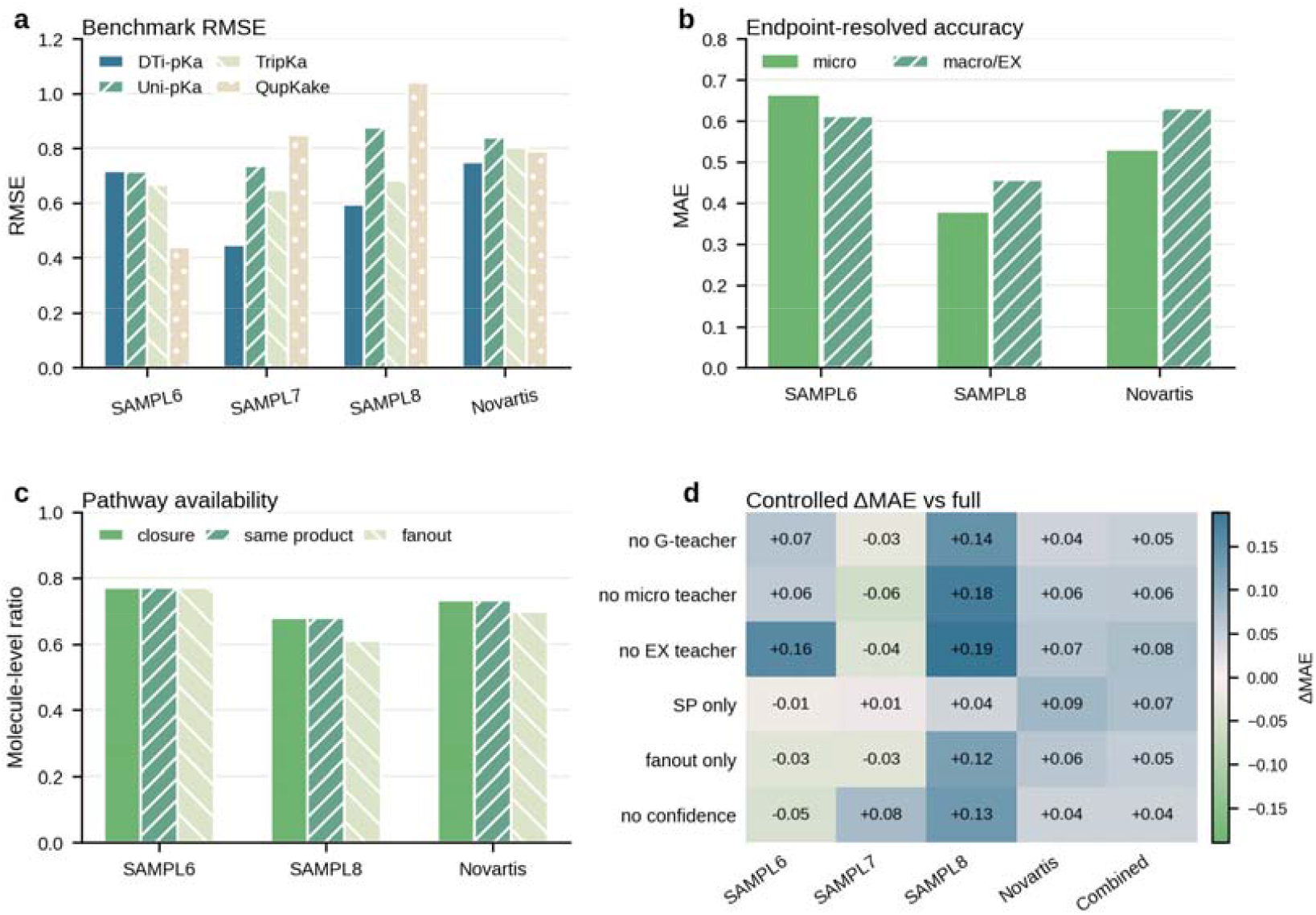
External benchmark controls and controlled ablation effects. **a**, Root-mean-square error (RMSE) of DTi-pKa and published comparator point estimates on four external benchmarks (SAMPL6, SAMPL7, SAMPL8, Novartis; n = 355 total). Published values are literature-context comparisons and carry no uncertainty; no controlled head-to-head evaluation was performed. **b**, Endpoint-resolved mean absolute error (MAE) by benchmark, shown separately for site-resolved (Micro) and macroscopic (EX) records. Bars are point estimates; sample sizes are reported in Supplementary Table 4. **c**, Fraction of molecules with available closure, same-product, and fanout pathways in external macroscopic (EX) datasets. Bars show molecule-level ratios for SAMPL6, SAMPL8, and Novartis; SAMPL7 is not shown because all its records are site-resolved (Micro). **d**, Paired ΔMAE of the six controlled ablation variants (A1–A6) relative to the full DTi-pKa model. Rows correspond to ablated components (defined in Supplementary Table 6), columns to benchmarks and the combined set. The color bar indicates ΔMAE in pKa units; positive values indicate higher error after component removal. Ablations are single-seed runs; intervals are unadjusted for multiple comparisons (Supplementary Information, Section S9).

**Fig. 5.**
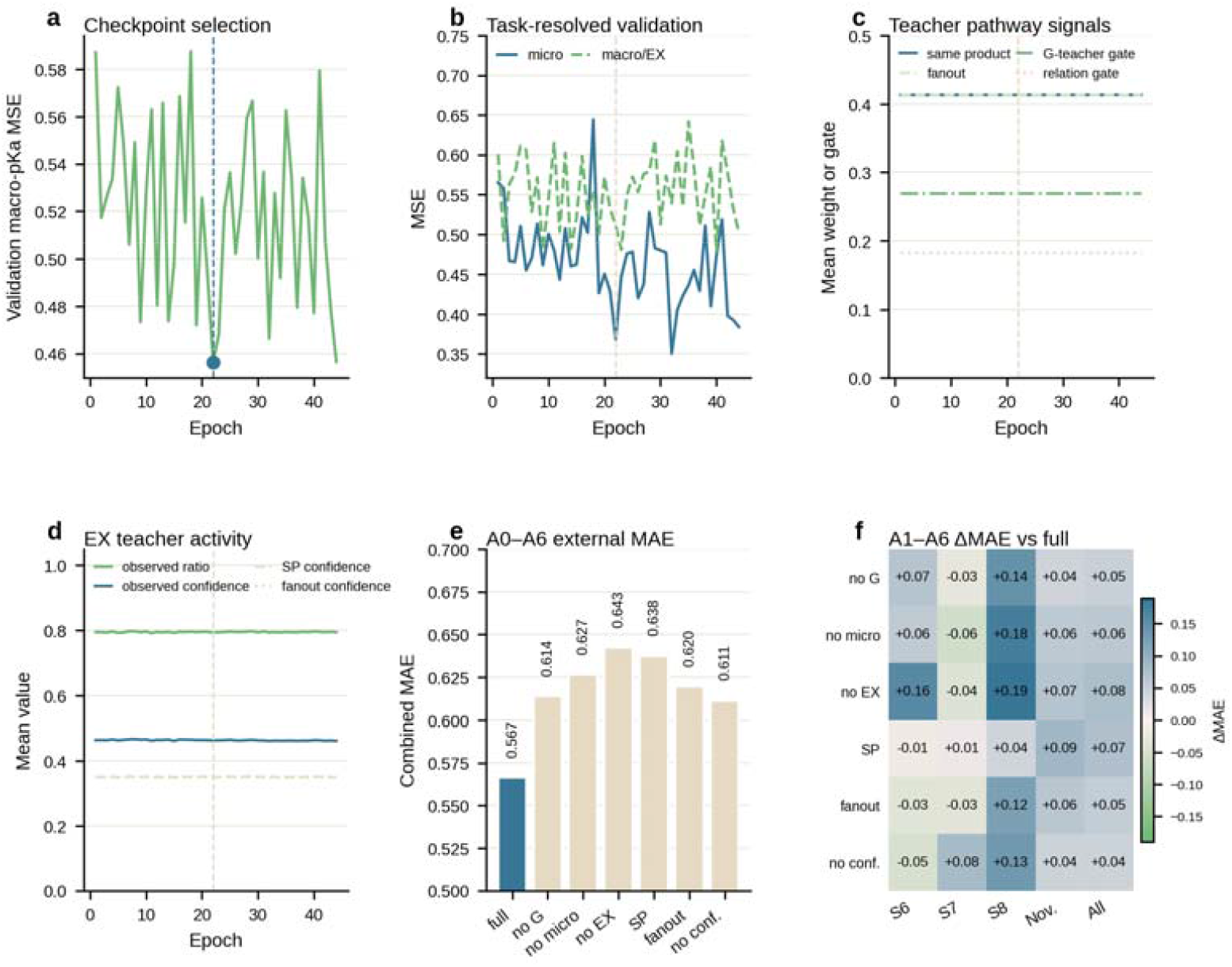
Validation diagnostics and controlled ablation results. **a**, Validation macroscopic pKa mean squared error (MSE) as a function of training epoch. The selected checkpoint (zero-based epoch 21, MSE 0.4563) is marked; selection was performed before any external benchmark analysis. **b**, Task-resolved validation error, shown separately for site-resolved (Micro) and macroscopic (EX) records. Bars are point estimates; sample sizes are reported in Supplementary Table 4. **c**, Teacher pathway signals during optimization, shown for same-product and fanout constraints. Curves represent the magnitude of the corresponding teacher loss terms; see Methods for definitions. **d**, Macroscopic (EX) teacher activity and effective confidence during training. Activity is defined as the teacher loss weight; effective confidence is the path confidence c_!JG_ (Methods). **e**, Pooled external mean absolute error (MAE) for the full model (A0) and the six controlled switch-off variants (A1–A6). Error bars represent 5% bootstrap confidence intervals computed from matched per-record absolute errors (10,000 resamples); ablations are single-seed runs and intervals are unadjusted for multiple comparisons (Supplementary Information, Section S9). **f**, Dataset-resolved ΔMAE for A1–A6 relative to the full model (A0). Rows correspond to ablated components (defined in Supplementary Table 6); columns correspond to the four external benchmarks and the combined set. The color bar indicates ΔMAE in pKa units; positive values indicate higher error after component removal.

Removing the macroscopic (EX) teacher, which calibrates the aggregated output against experimental macro-pKa, costs more (+0.0760; 95% interval +0.0392 to +0.1127) than removing the microscopic teachers (+0.0603; +0.0251 to +0.0951). Because the macroscopic pKa is a population-weighted sum, microscopic label errors partially average out during aggregation. The macro teacher instead calibrates the aggregate that is ultimately reported directly. Macroscopic consistency therefore matters more for final accuracy than microscopic numerical precision. This statement has direct implications for any physics-constrained predictor: constraint placement at the level of the reported quantity outperforms placement at the level of intermediate labels. The same logic explains why dropping the fanout direction costs more (+0.0711) than dropping the same-product direction (+0.0530). The fanout teacher distributes the correction across all cached product states, and it is the route through which macroscopic information reaches the largest fraction of the state space. The matched A6 rerun adds one protocol-level observation: confidence modulation was associated with a 0.0449 pKa unit lower pooled MAE (95% interval, 0.0142 to 0.0767). We also record an inversion between the selection metric and the external outcome. The A6 rerun reached a better best-validation MSE (0.4020) than V50 (0.4563) yet a worse pooled external MAE (0.6114 versus 0.5665). This cautions against reading the validation metric as a surrogate for external performance.

### Numerical closure of the identity remains an open frontier

We next examined whether the trained model satisfies the thermodynamic identity, not merely during training but at inference. In the recorded configuration, both the closure teacher and the closure forward scales are assigned zero weight. Numerical closure was therefore not enforced during training; the residuals below reflect this design choice and a model limitation. A CPU diagnostic on one Micro example and three selected EX examples measured same-product residuals of 0.711, 4.257, 0.130, and 1.522 pKa units, respectively. Fanout residuals were 3.762, 0.130, and 0.669 pKa units (Supplementary Information, Section S10). These residuals are heterogeneous, with some exceeding 4 pKa units, and are illustrative of selected cases rather than a comprehensive closure assessment.

The identity in equation (4) is algebraically exact for the retained state and path set. However, the agreement between the two learned heads is an empirical quantity, and the current checkpoint does not achieve numerical closure for all tested pathways. The thermodynamic character of the present model is therefore supervisory: teacher terms provide thermodynamic targets, while head-level numerical closure remains an open frontier that can be measured. Closing this gap—by activating the closure term at a nonzero weight—is a concrete next step.

### Full aggregation versus dominant-state baselines on site-resolved outputs

The same prediction that yields the macroscopic pKa also provides site-resolved microscopic pKa values and the underlying protonated-state populations. We selected 15 records with state-level predictions from the high-error region of the external sets. On this diagnostic subset, full aggregation gave a mean absolute error (MAE) of 1.5548 pKa units and a root-mean-square error (RMSE) of 1.7168 pKa units. A dominant-state-only baseline, which uses only the most populated protonated microstate for aggregation, gave an MAE of 1.5823 and an RMSE of 1.7296 (Supplementary Fig. 6 and Supplementary Information, Section S10). The subset MAE is 2.7 times the pooled value (0.5665), confirming that it is a diagnostic selection rather than a representative sample.

The mean per-record difference is 0.0275 pKa units in favor of full aggregation. The median per-record difference is zero, indicating that the two methods perform equally on most records; the baseline is better on 4 of 12 records, and the largest difference (0.3200, Novartis1) arises from a single record. The median path-selection sensitivity is 1.2756 pKa units and the maximum is 3.2604, where path-selection sensitivity is defined as the range of predicted macroscopic pKa across tested path selections. These results define the limits of the site-resolved output: on this diagnostic subset, the dominant state alone was not always sufficient, and path selection could materially affect the reported macroscopic pKa. The benefit of full aggregation is concentrated in records where the population is fragmented, suggesting that full aggregation is most valuable when multiple protonated microstates contribute comparably to the equilibrium.

### Failures localize the new bottleneck: protonated-microstate enumeration

Across the combined external set, 84.3% of Novartis predictions fell within 1 pKa unit of experiment and 99.3% within 2 units; SAMPL6 carried the widest error tail (Fig. 6). To convert this distribution into mechanistic understanding, we dissected the worst predictions. Novartis_134_ (absolute error 2.790 pKa units) enumerated 14 isomers, retained 11, recorded 26 cycle paths and carried a dominant retained-state population of 99.72%, so its error is not a fragmentation problem. Novartis_6_ (error 1.936) retained 10 states with a dominant population of only 45.00%. SAMPL8_3_ (error 1.848) retained 4 states with a dominant population of 51.39%. Both cases place comparable weight on multiple retained states, and the reported macro value inherits that fragmentation. State retention is governed by a model-computed absolute free-energy threshold of 15.0, so the retained set is itself a model output.

**Fig. 6.**
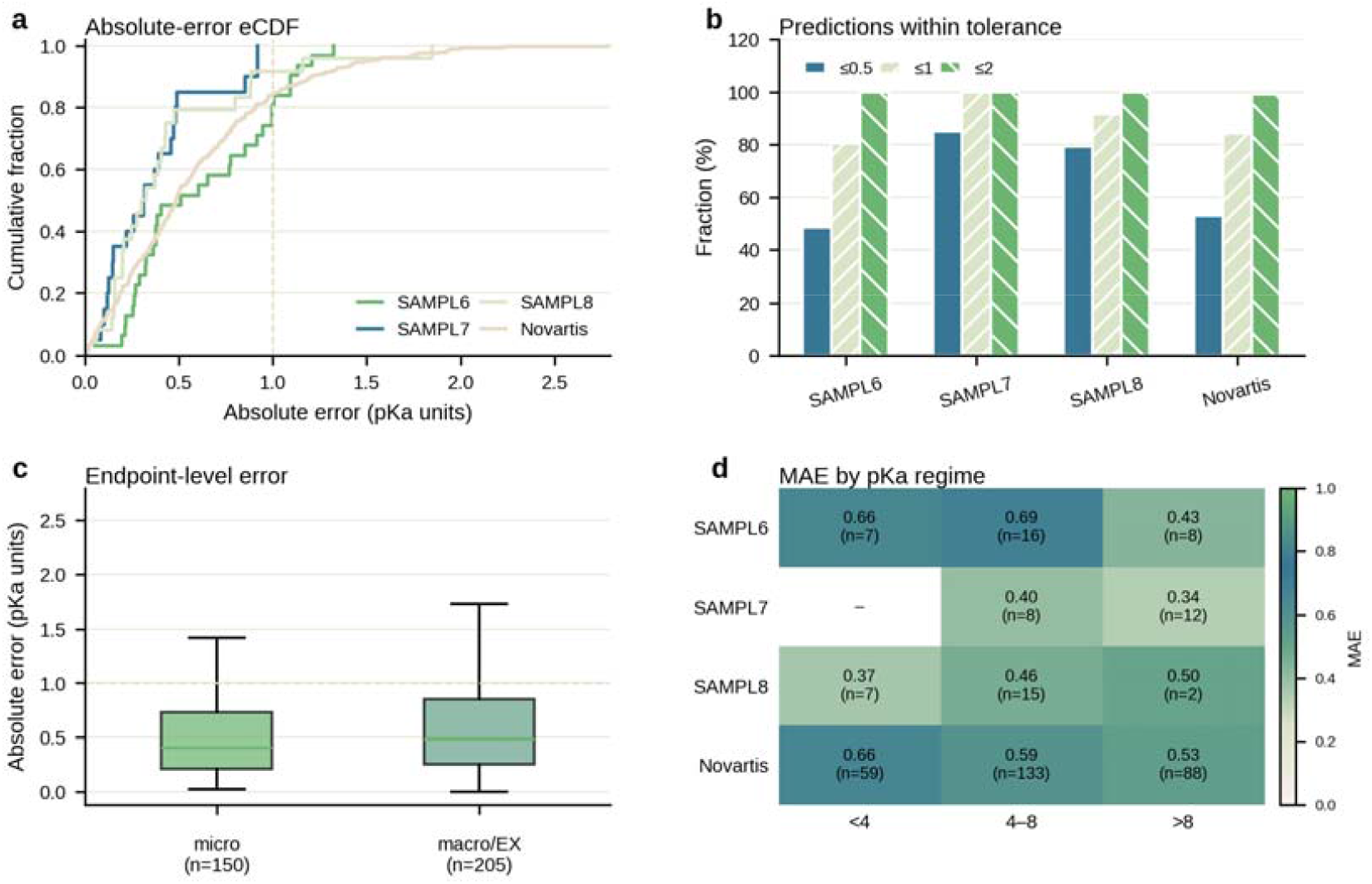
External error landscape of DTi-pKa. **a**, Empirical cumulative distribution of absolute error (pKa units) for each external benchmark. Curves show the fraction of predictions below a given absolute error. **b**, Fractions of predictions within 0.5, 1, and 2 pKa units, grouped by benchmark. Bars are point estimates; sample sizes are reported in Supplementary Table 4. **c**, Endpoint-level absolute-error distributions for site-resolved (Micro; n = 150) and macroscopic (EX; n = 205) records. Boxes show the median and interquartile range, whiskers extend to 1.5× the interquartile range, and points denote outliers. **d**, Mean absolute error (MAE) stratified by benchmark and experimental pKa regime (<4, 4–8, >8). The color bar indicates MAE in pKa units; numbers in cells denote sample sizes. Blank cells indicate no records in that stratum.

Whether the largest errors arise from enumeration gaps, representation limits, or label noise is not established here. These diagnostics identify targets for manual chemical review rather than assigning failure mechanisms, and each hypothesis remains a testable follow-up analysis. The largest errors concentrate where the protonated-microstate ensemble is fragmented, identifying enumeration completeness as the new rate-limiting step—consistent with the exactness conditions of equation (4), which require the relevant protonated microstates to be included. An adjudicated enumeration rule set and a larger state-level export would test these hypotheses directly.

## Discussion

DTi-pKa reframes pKa prediction from regression on labels to regression on a thermodynamic identity. An exact identity removes the deprotonated ensemble algebraically, reducing the 2^N^ protonation-state enumeration barrier to O(n+E) aggregation over retained protonated microstates and unique first-dissociation paths. The dual-head graph neural network assigns each quantity its thermodynamic role—one head for molecular free energies, one for site- and path-resolved dissociation constants—and couples them through training constraints. Controlled ablations show that this supervision does measurable work; the ranking of degradations (macroscopic teacher above microscopic teachers, fanout above same-product) reflects how thermodynamic information flows through the model. The broadest methodological finding is that constraint placement at the level of the reported quantity, the macroscopic equilibrium, matters more than microscopic label precision. This identifies a transferable design principle for physics-informed networks: constrain the quantity that is ultimately reported^2,4,34^. The principle reaches beyond pKa prediction to any domain where a governing identity can replace an ensemble enumeration.

The framework occupies a quadrant left empty by existing approaches. Black-box graph predictors are fast and accurate on familiar chemotypes, but they return no equilibrium and their microscopic and macroscopic outputs are not guaranteed to agree. Constant-pH molecular dynamics^23^ and free-energy methods such as Epik^24,25^ and MCCE2^26^ start from microscopic states and integrate their thermodynamics, but at hours per molecule, which confines them to candidate-level studies. QupKake^19^ and TripKa^20^ gain accuracy from quantum information at the cost of the same state-space and preprocessing burden that DTi-pKa avoids. DTi-pKa combines thermodynamically consistent aggregation at screening scale with the site-resolved interpretability of an ensemble method and the latency of a graph network.

The boundaries of the claim are specific. First, the identity eliminates the deprotonated ensemble, not the protonated one: prediction quality now depends on the completeness of the protonated-microstate ensemble generated by the SMARTS rules, and the failure cases above show where that dependence costs accuracy. Second, the model reads 2D molecular graphs without three-dimensional conformation or quantum descriptors; this is a deliberate trade of representation depth for screening scale, and the SAMPL6 deficit on high-polarity fused heteroaryls marks where that trade is most expensive. Third, the formalism covers first dissociation of organic molecules in water at room temperature; its form suggests, but does not prove, recursive extension to polyprotic systems. Each boundary is actionable: an adjudicated enumeration rule set addresses the first, uncertainty calibrated from the model’s own clarity statistics addresses the second, and recursive aggregation addresses the third.

For the wider community, the work demonstrates a transferable design principle: embed an identity that eliminates an intractable computation, enforce it as a soft constraint, and use ablations and closure diagnostics to test whether the constraint is causal rather than merely regularizing^1,2,5^. Three experiments remain explicitly open: a training run with the closure term active at a nonzero weight, matched-budget multi-seed ablations, and a single-head end-to-end network of matched capacity. Together these would separate the contributions of architecture, supervision, and seeds. A prospective, prediction-first drug-design demonstration would complete the translation of the site-resolved output into practice. The site-resolved, population-aware output provides a physical foundation for downstream ADMET modeling—solubility, logD, permeability—and for prospective molecular design at library scale^14^.

## Conclusion

We have shown that an exact thermodynamic identity can replace the exponential enumeration of deprotonated states in macroscopic pKa prediction. By expressing the macroscopic first-dissociation constant as a population-weighted sum over protonated microstates and unique first-dissociation paths, the identity reduces aggregation to O(n+E) and removes the 2^N^ barrier that has separated rigorous ensemble methods from screening-scale use. Implemented in DTi-pKa, a dual-head graph neural network trained with thermodynamic constraints, this framework achieves competitive external accuracy while returning site-resolved micro-pKa values, protonation-state populations, and path-level diagnostics.

Controlled ablations demonstrate that thermodynamic supervision does measurable work, and that constraint placement at the reported macroscopic equilibrium improves generalization more than microscopic label precision—a transferable design principle for physics-informed machine learning beyond pKa prediction. The generalization audit and closure diagnostics further show that the remaining gap is measurable and localized: numerical head-level closure and protonated-microstate enumeration completeness are the open frontiers.

Future work will extend first-dissociation aggregation to polyprotic systems, integrate uncertainty quantification to flag predictions outside the training chemical space, and complete closure-active training and single-head controls to separate the contributions of architecture, supervision, and seeds. The site-resolved, population-aware output provides a physical foundation for downstream ADMET modeling—solubility, logD, permeability—and for prospective molecular design at library scale.

## Methods

### Thermodynamic framework

The macroscopic first-dissociation constant is defined by equation (1) under the ideal-solution standard-state convention with dimensionless activities; the standard state fixes a(H^+^) = 1 without prescribing a physical proton concentration. Free energies and microscopic constants are defined by equation (2); the sign convention is fixed by *Ka*_*i,alpha*_ *= exp(ell*_*i,alpha*_) and *pKa*_*i,alpha*_ *= -ell*_*i,alpha*_*/ln 10*. The interconversion relation *Ka*_*ik*_ *= exp(g*_*i*_ *-g*_*k*_) follows from the common standard state and yields the population weights of equation (3). Unique-product bookkeeping assigns each distinct first-dissociation product to one representative parent-and-site pair, and the central identity (equations (4, 5)) follows by substitution; its exactness conditions (dimensionless activities, inclusion of the relevant protonated microstates and distinct products, consistency among convergent paths, canonical products counted once) are stated in the Results and derived in the Supporting Information. The log-sum-exp form (equations (6, 7)) is used for stable evaluation with maximum subtraction. The aggregation complexity O(n+E) applies to aggregation over the retained state and path set, not to the upstream generation of protomers and tautomers.

### Protonation-state enumeration and graph construction

The executable SMARTS catalogue^35^ contains 205 entries (94 acidic and 111 basic labels, 202 unique patterns); the catalogue file lists the rule index, the SMARTS pattern, the titration-site atom map number and the acidic or basic label. Several entries are deliberately broad, including one that matches any neutral trivalent nitrogen, so the catalogue admits chemically permissive first-dissociation paths; formal precision and recall would require a manually adjudicated ionizable-atom reference set, which we do not yet have (Supplementary Table S3). The enumeration pipeline normalizes the input, detects candidate acidic, basic and amphoteric sites, generates intramolecular proton-transfer candidates, enumerates tautomers through SMIRKS transformations with an RDKit fallback^36^, canonicalizes and deduplicates structures^37^, and retains protonated states carrying acidic or amphoteric sites. State retention uses a model-computed absolute free-energy threshold of 15.0, so the retained set is itself a checkpoint-dependent model output (Supplementary Figs. S1 and S2). The pipeline does not generate all 2^N^ protonation patterns and then filter them; only chemically reachable candidates are formed. First-dissociation product graphs and canonical product keys are generated for path bookkeeping and teacher diagnostics; products never enter the macroscopic sum as independent free-energy terms. Inputs are used as supplied; 216 of the 355 benchmark records carry a net formal charge. Graphs use 49-dimensional node features and 6-dimensional edge features, a six-layer message-passing encoder^38^ with hidden dimension 300 and dropout 0.2, pair features with k = 2 over at most 24 neighbours, and an H^+^ anchor weight of 0.01 (Supplementary Table S1). The checkpoint state dictionary contains 14,468,923 tensor elements; the training logs do not record a separate trainable-parameter inventory.

### Dual-head architecture and thermodynamic supervision

The G-head predicts a dimensionless molecular free-energy scalar. The dissociation head predicts *ell = ln Ka* for a parent state and a specific deprotonation path, with *pKa = -ell/ln 10*; in the machine-readable exports this scalar is stored under the field name ΔG_micro over RT_, which equals ell. For the same-product teacher, G_teacher_(HA) = G(A^-^) + ell; for the fanout teacher, G_teacher_(A^-^) = G(HA) - ell. Teacher strength is modulated by the clarity of the dissociation path and of the anchor representations.

Clarity is defined explicitly: for a detached high-dimensional G representation z_G_ in dimension d, c_G_ = tanh((||z_G_ ||/sqrt(d))/2); for each dissociation path, attention diagnostics are combined as

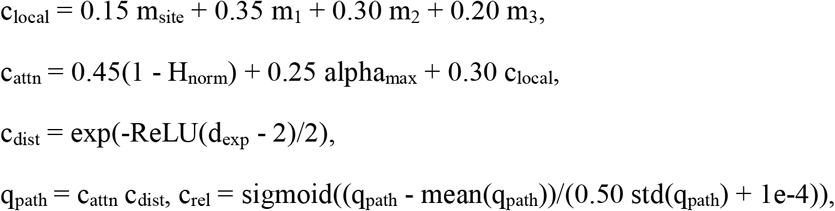

b_local_ = sigmoid(8(c_local_ - 0.18)) and c_ΔG_= 0.05 + 0.95(0.45 q_path_ + 0.35 c_rel_ + 0.20 b_local_), with a micro label-fit term c_label_ = exp(-|ΔG_pred_ + ln(10) pKa_exp_|/2.5). A PathConfidence multilayer perceptron (4 or 5 to 32 to 16 to 1, SiLU, sigmoid output) fuses these quantities into the path confidence that scales the teacher loss; no hard minimum-confidence gate is applied. The recorded configuration assigns nonzero weights to the G teacher (0.08) and the soft-ΔG teacher (0.015), and zero weight to the closure teacher with all three closure forward scales set to zero. The recorded configuration assigns nonzero weights to the G teacher (0.08) and the soft-ΔG teacher (0.015), and zero weight to the closure teacher, with all three closure forward scales set to zero; closure residuals for this checkpoint are reported under “Learned heads do not yet numerically close the identity”.

### Training and evaluation

The transfer-learning set integrates experimental pKa measurements from iBonD^39^, ChEMBL^40,41^ and DataWarrior-derived files^42^ (raw-file and retained-cache counts in Supplementary Information S6). The split uses a 0.20 validation ratio with seed 7. Training used AdamW (learning rate 6e-6, weight decay 2e-4), bf16 automatic mixed precision, micro batch size 32, external gradient accumulation up to 8, and a three-phase curriculum of 14/14/16 epochs (44 logged of 50 configured; the run was executed in single-epoch segments with resume and the optimizer state was not restored) (Supplementary Table S2). The V50 reference used min_epochs = 8; the A6 rerun used min_epochs = 44. The publication checkpoint was selected by validation macro-pKa MSE (0.4563212141) at zero-based epoch 21 before any external benchmark analysis; SAMPL6, SAMPL7, SAMPL8 and Novartis were then evaluated once with the fixed checkpoint, with no failed predictions (355/355 valid). The separate A6 no-confidence rerun used the same 44 logged epochs, selected zero-based epoch 40 (MSE 0.4019948637), and was evaluated with its own checkpoint. Records carry one of two label types: site-resolved (Micro) records, which carry a populated acidic_idx, and macroscopic (EX) records, which do not; the two types are different estimands and are reported separately. Paired delta-MAE values and 95% bootstrap intervals were computed on matched per-record absolute errors (10,000 resamples, seed 20260725, unadjusted for multiple comparisons). Standard errors of the per-benchmark MAE were computed over per-record absolute errors. The cache-overlap audit compares canonical SMILES against the fine-tuning cache; the pretraining corpus was not included. Inference timing was measured on an NVIDIA GeForce RTX 4080 with one warm-up inference excluded. Main-text values are rounded to four decimal places; full-precision values and per-record predictions are provided in the Supplementary Information and the machine-readable source-data files (Supplementary Information S12).

## Data availability

The SAMPL6, SAMPL7 and SAMPL8 benchmark sets are available from the SAMPL challenge publications and repositories. Per-record predictions for all external benchmarks are provided as machine-readable source data with this paper (Supplementary Information S12). The training data derive from public sources including iBonD, ChEMBL and DataWarrior-derived files.

## Code availability

The benchmark and evaluation source code are freely available at https://github.com/IDLab2026/DTi-pka.

## Supporting Information

Additional methodological details, extended derivations, dataset statistics, complete ablation results, per-molecule predictions and reproducibility information are provided in the Supporting Information.

## Acknowledgements

This work was supported by the National Foreign Expert Program (Grant No. H20250869), the Fundamental and Interdisciplinary Disciplines Breakthrough Plan of the Ministry of Education of China (Grant No. JYB2025XDXM609), the National Key Research and Development Program of China (Grant No. 2022YFC2804900), the National Natural Science Foundation of China (Grant Nos. 22277006, 92259302 and 22403069), the Beijing Natural Science Foundation (Grant No. 7242195), the National Talent Program (Grant Nos. CZ000935, KZ667301 and KZ0139), National Science and Technology Major Project for Innovative Drug Research and Development (Grant No. 2026ZD1807000) and the 2026 Special Project for Education Development (Third Batch) – Basic Scientific Research Operating Funds for Higher Education Institutions (Grant No. CZ000713).

